## Supplementary Material for "Recurring adaptive introgression of a supergene variant that determines social organization"

**This PDF file includes:**

Supplementary Notes 1 to 10

Supplementary Figs. 1 to 16

### Supplementary Notes

#### Supplementary Note 1: Dating divergence times of *Solenopsis* species and the supergene

*Background.* The age of the supergene can be estimated by building a dating phylogeny calibrated using nodes retrieved from fossil-calibrated trees. The supergene was first dated to be at least 0.39 million years old by comparing the synonymous substitution rate between SB and Sb to the synonymous substitution rate between two species of leafcutter ants, *Atta cephalotes* and *Acromyrmex echinatior*. This dating had been calibrated using an estimated divergence time of 10 million years between the leafcutter taxa^3^; recent divergence time estimates^4–7^ range from 8 to 17.5 million years ago (mya). A recent study of the fire ant system^8^ inferred the supergene to be an ancestral polymorphism originating at a node between the divergence of *S. saevissima* and the divergence of *S. interrupta*. This study estimated that the supergene is at least 0.506 million years old by building a phylogeny with five nuclear genes and *Solenopsis xyloni* as an outgroup. However, this study assumed that *S. xyloni* diverged from the other *Solenopsis* species 4.5 mya, despite citing a study^5^ that estimated that divergence between *S. xyloni* and *S. invicta* at 3.91 mya.

*Available calibration nodes*: To increase power and dating accuracy relative to previous studies, our approach was to build dated phylogenetic trees from single-copy genes from across the genome, and to calibrate divergence based on two nodes. The first calibration node separates *S. geminata* from the remaining species in the tree. Morphology and molecular data show that *S. geminata* is in the same branch as *Solenopsis xyloni^9–11^*. Therefore, we consider the divergence time between *S. geminata* and *S. invicta* to be identical to the reported divergence time between *S. xyloni* and *S. invicta* (3.91 mya^5^). For the second calibration node, we included the more anciently diverged species *Solenopsis* *fugax*, for which a genome sequence is available ([GCA_003595255.1](https://www.ncbi.nlm.nih.gov/assembly/GCA_003595255.1/)^12^). The divergence time between *S*. *fugax* and *S. xyloni* — and hence between *S*. *fugax* and the species in our study — has been estimated to be 25.2 million years ago^6^.

*Methods and results:* To avoid mapping-biases in species distant to *S. invicta*, we constructed a short-read based genome assembly for each sample in the dated phylogeny analysis using Masurca^13^ (v 3.3.7). Specifically, we generated assemblies for a number of individuals used in the main analysis: *S. richteri* SB and Sb (reads used: SRA accessions [SRR9008200](https://www.ncbi.nlm.nih.gov/sra/?term=SRR9008200), [SRR9008133](https://www.ncbi.nlm.nih.gov/sra/?term=SRR9008133)), *S. macdonaghi* SB and Sb ([SRR9008142](https://www.ncbi.nlm.nih.gov/sra/?term=SRR9008142), [SRR9008173](https://www.ncbi.nlm.nih.gov/sra/?term=SRR9008173)), *S. invicta* SB and Sb ([SRR9008253](https://www.ncbi.nlm.nih.gov/sra/?term=SRR9008253), [SRR9008228](https://www.ncbi.nlm.nih.gov/sra/?term=SRR9008228)), *S. AdRX* SB and Sb ([SRR9008232](https://www.ncbi.nlm.nih.gov/sra/?term=SRR9008232), [SRR9008168](https://www.ncbi.nlm.nih.gov/sra/?term=SRR9008168)), *S. interrupta* SB ([SRR9008217](https://www.ncbi.nlm.nih.gov/sra/?term=SRR9008217)), *S. megergates* SB ([SRR9008215](https://www.ncbi.nlm.nih.gov/sra/?term=SRR9008215)), *S. geminata* (gem1, see Supplementary Data 1), *S. pusillignis* (Lad4-1, see Supplementary Data 1), and *S. saevissima* (Par2-5, see Supplementary Data 1, [SRR9008150](https://www.ncbi.nlm.nih.gov/sra/?term=SRR9008150), [SRR9008158](https://www.ncbi.nlm.nih.gov/sra/?term=SRR9008158)). Additionally, we included multiple published *S. invicta* assemblies: [GCF_016802725.1](https://www.ncbi.nlm.nih.gov/assembly/GCF_016802725.1), [GCA_010367695.1](https://www.ncbi.nlm.nih.gov/assembly/GCA_010367695.1), [GCA_009299975.1](https://www.ncbi.nlm.nih.gov/assembly/GCA_009299975.1), [GCA_009299965.1](https://www.ncbi.nlm.nih.gov/assembly/GCA_009299965.1), [GCA_018691235.1](https://www.ncbi.nlm.nih.gov/assembly/GCA_018691235.1) (generated from brothers of the sample CGI1-1-bigB from the main analysis), and Sinv_gnGA^8,14–16^, as well as *S. fugax* ([GCA_003595255.1](https://www.ncbi.nlm.nih.gov/assembly/GCA_003595255.1)^12^). For each assembly, we identified complete and single copy conserved orthologs using BUSCO (v.4.05, i.e., the same version as used for the main analysis, Supplementary Data 2). The assembly statistics, sample details and BUSCO scores are summarized in Supplementary Data 4. The new genome assemblies can be downloaded from https://wurmlab.com/data/supergene_introgression/

Using the nucleotide sequences (coding sequence as reported by BUSCO) of those genes present in all assemblies (n = 22), we constructed per-gene alignments using prank^17^ (v170427) before trimming alignment gaps (phyx: pxclsq -p 1.0) and concatenation (phyx: pxcat) of those genes located on chromosomes 1 to 15 (*i.e.,* not on chromosome 16) in *S. invicta* (n = 2,161). Each gene was one partition and we used previously inferred best substitution models (Supplementary Data 2). We then used the concatenated alignment (length: 2,906,225 nucleotides) to construct a maximum likelihood tree with 1,000 ultrafast bootstraps, and phylogenetic dating with 100 replicates using IQ-TREE^18^ (v2.1.3) and the age calibrations mentioned above.

The resulting maximum likelihood tree topology is highly supported and congruent with the species tree shown in Fig. 1 (chromosomes 1–15) (Supplementary Fig. 2). As expected, individuals of the same species carrying different supergene variants always cluster together. Importantly, this phylogenetic inference also places *S. richteri* as sister to *S. invicta*/*S. macdonaghi*, and the clade with these two groups as sister to a clade composed of *S. megergates* and *S. AdRX*, congruent with the main species tree (Fig. 1). The divergence time between *S. invicta* and *S. saevissima* was inferred to be 1.76 mya, to *S. megergates + S. AdRX* 1.09 mya, and to *S. richteri* 1.01 mya (Supplementary Fig. 2). Considering *S. invicta* and *S. macdonaghi* as one species complex harboring the supergene (see main text) and, assuming that *S. richteri, S. megergates, S. interrupta* and *S. AdRX* acquired the supergene via introgression, the origin of the supergene is placed after the speciation event leading to *S. richteri* and *S. invicta/S. macdonaghi*, between 1.01 mya and 0.67 mya. These inferred divergence times are longer than previously reported^8^, likely due to the vastly larger dataset (2161 *vs.* 5 genes) and the additional age calibration of *S. fugax* (repeating the analysis using each of the two age calibrations independently resulted in similar inferred ages, data not shown).

As an additional analysis, we built a dated species tree containing all individuals in our study (Fig. 1), making use of the dense sampling to obtain more precise ages of the diversification of *S. invicta*. We used the node ages inferred above to date the divergence of the major species (nodes with branches leading to *S. geminata*: -3.91 mya, *S. pusillignis*: -2.67 mya, *S. saevissima*: -1.76 mya, *S. megergates+S. AdRX*: -1.09 mya and *S. richteri*: -1.01 mya; but not to *S. macdonaghi* due its polyphyly). We concatenated the alignments of the single-copy genes from chromosomes 1 to 15 used to produce the species tree (Fig. 1). We used IQ-TREE 2 as above to produce a dated phylogeny (Supplementary Fig. 3).

As expected, there is a high congruence between the dated tree produced in this analysis and the one produced above. This can be seen by looking at nodes present in both analyses, but which we did not restrict in the second analysis (Supplementary Figs. 2 and 3):

- diversification date of *S. saevissima* (1.26 mya and 1.43 mya, respectively in the first and second analysis)
- divergence between *S. interrupta* and the remaining crown species (1.20 mya and 1.23 mya, respectively in the first and second analysis)
- divergence between *S. megergates* and *S. AdRX* (0.97 mya and 1.06 mya, respectively in the first and second analysis)
- diversification of the *S. invicta*/*S. macdonaghi* (0.67 mya and 0.68 mya, respectively in the first and second analysis)

This second analysis shows that the *S. invicta*/*S. macdonaghi* lineage diversified 0.81 mya with a large radiation 0.68 mya. This suggests that the supergene arose between 1.01 mya and 0.81 mya.

To infer a more precise age of the origin of the supergene, we repeated the second analysis, using the genes mapped to the supergene region. For this analysis, we excluded the calibration of *S. richteri–S. invicta*. By doing this, we avoid biases from forcing node ages between groups of higher rates of introgression and the potential of SB-Sb recombination or gene conversion in the supergene region. The age of the split between *S. richteri* and *S. invicta/S. macdonaghi* was almost identical (1.06 mya instead of 1.01 mya). Based on this inference, the split between both supergene variants (SB and Sb) and hence the origin of the supergene occurred 0.97 mya (Supplementary Fig. 4). Congruent with the dated species tree (chromosomes 1 to 15) showing a radiation of *S. invicta/S. macdonaghi* 0.81 mya, the SB variant of the supergene diversified 0.82 mya, while the Sb variant of the supergene diversified 0.89 mya.

We used the dated supergene tree to estimate when introgression events occurred. The majority of introgression events from *S. invicta* into other species occurred between 0.89 mya and 0.75 mya (*S*. *richteri* 1: 0.89 mya; *S. richteri* 2: 0.85 mya; *S. megergates*: 0.85 mya; *S. AdRX*: 0.95 mya but the node has zero support, 0.85 mya based on its position in the concatenation tree (Supplementary Fig. 7) as sister to a clade consisting of AR186-1, AR186-3, AR187-1, AR187-5, SRR9008173_mac−146, SRR9008257_mac−147 in both the concatenation tree (Supplementary Fig. 7) and the supergene tree (Fig. 1e, right; *S. interrupta*: 0.75 mya), *i.e.*, in the first quarter of the supergene’s existence. The *S. megergates* (GCa3) introgression event was dated at 0.4 mya, reflecting allelic differences among a local population (0.34–0.44 mya between pairs of *S. richteri*, *S. macdonaghi* and *S. invicta* from the same location, Supplementary Fig. 2) but older than brothers (0.14–0.26 mya between *S. invicta* SB and Sb brothers, Supplementary Fig. 2).

#### Supplementary Note 2: Nucleotide diversity (π) of Sb in *S. invicta/macdonaghi* and each of the two introgressed clades of *S. richteri*

We aimed to compare the genetic diversity in each of the two clades of introgressed *S. richteri* Sb, with an *S. invicta/macdonaghi* clade. As representatives of the *S. invicta/macdonaghi* clade, we used the 62 haploid males that form the monophyletic group that is sister to the *S. richteri* clade 2. We used all 10 haploid males from *S. richteri* clade 1 (Fig. 1e), and all 34 *S. richteri* males from clade 2. To avoid potentially ambiguous regions of the genome, we only considered SNPs in the vicinity of single-copy gene regions, defined as the exons, introns and 1000 bp upstream and downstream of each of the 210 single-copy “BUSCO'' genes in the supergene region of chromosome 16. After collapsing overlapping regions, this provided 141 regions covering 2.16 Mbp. We measured nucleotide diversity π using the ‘nucleotide.diversity.within’ function in the R library PopGenome^2^, summed across all 141 regions and divided by their combined total length of 2.16 Mbp (Supplementary Fig. 5). The nucleotide diversities of the two *S. richteri* clades (0.0010 for clade 1 and 0.0015 for clade 2) were, respectively, 68% and 99% that of *S*. *invicta/macdonaghi* Sb (0.0015), suggesting that both introgressed clades harbor extensive diversity. This is consistent with substantial amounts of time having passed since introgression into *S. richteri*.

#### Supplementary Note 3: Recent introgression of Sb into *S. megergates*

One of the two introgressed samples of *S. megergates* (introgression 3 in Fig. 1e), groups inside a clade of three *S. invicta*/*macdonaghi* Sb samples collected within 5.2 km of each other. If the Sb haplotypes carried by these samples were particularly similar, this would suggest that this introgression happened recently.

To test this, we measured the Bray-Curtis dissimilarity index for all 9,591 possible pairs of the 139 Sb haplotypes. For this we applied the R package vegan^1^ using genotypes of the 24,435 SNPs that were bi-allelic among the Sb samples and mapped to the supergene (Supplementary Data 3). The distribution of dissimilarity values is shown in Supplementary Fig. 6. We note the following points:

1. The pairwise dissimilarity values between the single *S. megergates* male (introgression 3) and each of the three *S. invicta/macdonaghi* Sb males in the clade were ~0.22.
2. These values are lower than the average pairwise dissimilarity values between the *S. richteri* samples in introgression 1 and the two closest *S. invicta/macdonaghi* samples (respectively 0.60 and 0.62).
3. These values are also lower than average pairwise dissimilarity values between the *S. richteri* samples in introgression 2 and the four closest *S. invicta/macdonaghi* samples (ranges between 0.55 and 0.59).
4. In contrast, these values are within the range of dissimilarities measured between Sb haplotypes of samples collected from one colony (ranges between ~0 and 0.29).

We conclude that the introgression of this Sb haplotype into *S. megergates* (introgression 3) occurred more recently than either of the two introgressions of Sb into *S. richteri*. The greater age of the *S. richteri* introgressions is consistent with the genetic diversity seen within both introgressed clades (Supplementary Note 2).

#### Supplementary Note 4: Phylogenetic analysis of *Solenopsis* using concatenated SNPs of single-copy genes

As an alternative to the coalescent-based tree reconstruction approach used in the main text, we built phylogenetic trees using a gene concatenation approach similar to the method used in a previous phylogenetic study of *Solenopsis* fire ants^8^. We retrieved SNPs located in the coding regions of the single-copy “BUSCO'' genes using bedtools^19^. We created multiple sequence alignments using vcf2phylip^20^, concatenated these alignments with the phyx^21^ tool pxcat, and generated two supermatrices. The supermatrix of chromosomes 1-15 included 428,888 sites from 5,450 single-copy genes and the supermatrix of the supergene region of chromosome 16 included 20,067 sites from 210 single-copy genes. We built a phylogenetic tree from each supermatrix using RAxML-NG^22^ (v0.9.0) with a single model (GTR + G, unpartitioned) and 10 independent searches (each with two starting trees, one parsimony and one random), evaluated support with 100 bootstrap replicates and computed transfer bootstrap expectation^23^ values.

The results obtained (Supplementary Fig. 7) were consistent with those from the coalescent-based tree reconstructions (Fig. 1e). As in the coalescent-based tree, there was unambiguous support for one introgression of Sb from *S. invicta/macdonaghi* into *S. interrupta*, two into *S. megergates* and one into *S. AdRX*. Although topologies from both tree-building approaches also indicated that Sb introgressed from *S. invicta* into *S. richteri,* a notable difference between results from the two approaches was that the *S. richteri* Sb haplotypes are placed into one monophyletic group in the concatenated alignment tree (Supplementary Fig. 7) and into two monophyletic groups in the coalescent-based tree. Although we cannot rule out the possibility that Sb introgressed only once into *S. richteri*, we note that the monophyletic grouping of Sb in the concatenated alignment trees could plausibly result from recombination between separately introgressed Sb haplotypes within the *S. richteri* population. Such recombination could take place within *Sb/Sb* queens, which occur in *S. richteri* more frequently than in *S. invicta^24^*.

#### Supplementary Note 5: Quantifying introgression using branch lengths of gene trees using QuIBL

Incongruence between gene trees and the species tree can be caused by introgression. However, it can also be caused by the differential retention of ancestral alleles in different lineages, a process known as incomplete lineage sorting (ILS). The method QuIBL^25^ (Quantifying Introgression via Branch Lengths; git commit e79a390990183f2804c2d5ca3feb014c2bbf5f04) allows distinguishing between ILS and introgression based on the distribution of internal branch lengths of triplet subtrees. QuIBL infers introgression when a bimodal distribution of internal branch lengths fits the data better than an exponential distribution.

We applied QuIBL to further test for introgression in the evolutionary history of *Solenopsis* using trees inferred from single-copy BUSCO genes located in the supergene region. We first focused on the inferred introgression even from *S. invicta* to *S. richteri.* Because QuIBL is more appropriate for small numbers of samples, as it examines every possible triplet in a dataset, we subsampled our dataset to 20 *S. invicta* samples, 20 *S. richteri* samples, 7 *S. saevissima* samples and 2 *S. pusilignis* samples (Supplementary Data 1), the two latter species having no evidence for an introgression event. We inferred maximum-likelihood gene trees using RAxML-NG^2^ (v0.9.0) with the model GTR + G and 20 starting trees (10 parsimony, 10 random), producing 107 gene trees. We used the phyx^3^ tool pxrr to root all gene trees with the outgroup *S. geminata* (sample “gem-1-bigB-m-majorityallele”). We detected introgressed loci using QuIBL with a total of 50 expectation–maximization (EM) steps and analyzed the results using the R package quiblR (https://github.com/nbedelman/quiblR). We distinguished between the ILS + introgression and ILS-only models based on the difference between Bayesian information criterion (BIC) scores, with a significance threshold equal to ∆BIC > 10. We used quiblR’s function “getIntrogressionSummary” to calculate the average introgression fraction for each pair of samples.

Our results indicated higher levels of introgression between Sb samples in *S. invicta* and in *S. richteri* relative to comparisons between SB samples in *S. invicta* and in *S. richteri*, as seen by high values of average introgression fraction in Supplementary Fig. 8. Furthermore, we examined the distributions of internal branch lengths for significant triplets involved in introgression events. The branch length distributions usually revealed two peaks (consistent with introgression) instead of an exponential decay (ILS only) in the comparison between Sb samples in *S. invicta* and in *S. richteri* (Supplementary Fig. 9).

We repeated this analysis to evaluate the introgression between *S. invicta* and the closely related species *S. megergates*, *S.* *AdRX* and *S. interrupta*. We used a subset of 17 samples representing all seven species in our dataset (Supplementary Data 1). This analysis also showed high values of average introgression fraction for the interspecific comparisons of Sb samples, but not for those of SB samples (Supplementary Fig. 10), consistent with introgression. Moreover, the branch length distributions usually revealed two peaks (introgression) instead of an exponential decay (ILS-only) in the interspecific of Sb samples (Supplementary Fig. 11).

#### Supplementary Note 6: Alleles shared between Sb and the SB of each species

Under a scenario with no introgression (with the supergene as an ancestral polymorphism originating before the speciation), patterns of allele sharing between the SB and Sb variants of different species would be explained by incomplete lineage sorting alone. If we take *S. invicta/macdonaghi* and *S. richteri* as an example, we have the following tree topology:

((richteri_Sb, invicta_Sb),(invicta_SB,richteri_SB)),saevissima)

We would expect the Sb variants to have the *same number of alleles* in common with the SB variant of *S. invicta* as with the SB variant of *S. richteri*. If we represent this using syntax similar to that used in the calculation of *d*-statistics, the number of BBBAA sites would be similar to the number of BBABA sites. This is illustrated in Supplementary Fig. 12a.

On the other hand, under the introgression scenario, the Sb variants share a more recent common ancestor with the SB of *S. invicta/macdonaghi* than with the SB of other species:

((richteri_Sb, invicta_Sb),invicta_SB),richteri_SB)),saevissima)

Under this scenario of introgression, we would expect that *Sb has more alleles in common* with the SB of *S. invicta/macdonaghi* than with the SB variant of *S. richteri*. The number of BBBAA sites would be greater than the number of BBABA sites (Supplementary Fig. 12a).

To test this, we focused on *S. invicta/macdonaghi* and *S. richteri*, for which we have sufficient samples carrying the Sb variant. We selected four groups of samples: SB and Sb of *S. invicta/macdonaghi* and SB and Sb of *S. richteri*. To limit any potential power biases, we retained equal numbers of samples from each group, while randomly selecting among those samples with fewer than 5% missing genotypes. *S. richteri* SB had the lowest number of samples available. Thus, we retained 28 samples from each of the four groups (Supplementary Data 1). We also included 11 samples of *S. saevissima*, a species with no introgression that we considered as an outgroup for this analysis (Supplementary Data 1).

We subset our dataset to the bi-allelic SNPs that map to the supergene. We kept only those SNPs that were invariant in the outgroup and that were genotyped in at least 22 of the 28 individuals in each of the four groups. For each SNP site, we classed the allele carried by the *S. saevissima* outgroup samples as ancestral, and the other allele as derived. We then counted the number of sites with the derived allele present in at least one of the Sb groups and also present in either the *S. invicta/macdonaghi* SB group or in the *S. richteri* SB group. These steps were done using the VariantAnnotation library^26^ in R. We found that Sb had 1,921 sites with a derived allele in common with *S. invicta/macdonaghi* SB samples and 994 alleles in common with *S. richteri* SB samples (*i.e.*, 1.93 times more Sb alleles shared with *S. invicta/macdonaghi* SB; exact binomial test, *p* < 10^-66^, 95 % confidence interval of 1.79–2.09 times). This difference is not expected under the scenario with no introgression, where the shared alleles are the result of incomplete lineage sorting. Instead, it supports the introgression scenario, with a more recent ancestor between Sb and the SB variant of *S. invicta/macdonaghi*. To ensure that these results are not driven by parallel selective pressures or mutational processes, we further subset the SNPs to those with a predicted synonymous effect (likely to be neutral), using the GenomicFeatures and VariantAnnotation (‘predictCoding’ function) libraries in R. This revealed that Sb had 222 alleles in common with *S. invicta/macdonaghi* SB samples and 92 alleles in common with *S. richteri* SB samples (*i.e.,* 2.41 times more Sb alleles were shared with *S. invicta/macdonaghi* SB; Exact binomial test, *p* < 10^-12^, 95 % confidence interval of 1.88–3.11 times).

To corroborate these results, we examined whether a similar pattern is seen for the other three species with evidence for introgression. Given our low sample size for these species, we selected a pair of individuals (one SB and one Sb) for each of the four species with putative introgression, as well as for *S. invicta*. The samples selected are:

- *S. AdRX*: SRR9008232_AdRX-134-Bra-bigB and SRR9008168_AdRX-135-Pos-littleb
- *S. richteri:* SRR9008197_ric-96-Ros-bigB and SRR9008136_ric-74-Ros-littleb
- *S. megergates*: SRR9008214_meg-130-BZ-bigB and SRR9008167_meg-127-BZ-littleb
- *S. interrupta*: SRR9008217_int-122-Car-bigB and SRR9008166_int-125-Car-littleb
- *S. invicta*: SRR9008275_inv-223-Cor-bigB and SRR9008163_inv-225-Mis-littleb

We also selected a single *S. saevissima* individual (SRR9008158_sae-47-Bel-bigB), representing a non-introgressed outgroup. For each of the four “focal” introgressed species, we considered a tree comparing that species with *S. invicta* and the outgroup:

((focal_Sb, invicta_Sb), invicta_SB), focal_SB)), outgroup)

For each of the four “focal” introgressed species, we counted how many derived alleles were present in both Sb samples (*S. invicta/macdonaghi* and the focal species) and either the SB of *S. invicta* or the SB of the focal species (*i.e.,* found in all three individuals; using a syntax similar to that used in the calculation of *d-*statistics, this comparison would be BBBAA *versus* BBABA, given the tree above). This was done using the script fasta2dfoil.py from the dfoil software^27^ (https://github.com/jbpease/dfoil , commit 3a69e2306a9f66d562cd3a0aaa08389d7fdbcbf1). Most genes had only a few such alleles (Supplementary Fig. 12b). Nevertheless, on average *S. invicta* had more private alleles in common with Sb than each of the focal species (two-tail paired t-tests, Bonferroni-corrected *p* < 0.05, test parameters in Supplementary Fig. 12b).

In summary, we found that SB shared more alleles with the Sb branch in *S. invicta* than in any of the other species. This pattern is consistent with the topology of the supergene tree, locating all Sb samples as a sister clade to the *S. invicta* SB samples (Fig. 1). These findings therefore further support the origin of Sb from the ancestral supergene region of *S. invicta* (which became SB), and then introgressing into each of the other species.

#### Supplementary Note 7: Phylogenetic weighting analysis of non-overlapping windows: Twisst comparison of alternative topologies along the supergene

To explore whether the phylogenetic relationship between samples varies across the supergene region, we used the Twisst^28^ method of phylogenetic topology weighting (<https://github.com/simonhmartin/twisst>, commit 4e4700120fc4f24e4cc24bd6ce446b347fae5e70). We focused on the putative introgression between *S. invicta/macdonaghi* and *S. richteri*, for which we have a large enough number of samples carrying the Sb supergene. We also included samples from species with no introgression: *S. pusillignis*, *S. saevissima* and *S. geminata*. To minimize reference bias (the reference assembly is based on a SB *S. invicta/macdonaghi* sample) we removed samples with more than 5% of missing genotypes (see Supplementary Data 1 for the samples used in this analysis). Using previously defined chromosomal locations of the assembly scaffolds^14^, we ordered the single-copy genes mapped to chromosomes 1 and 16 according to chromosomal position. We removed the four genes in scaffold NW_011795620.1 (scaffold length: 65 kb) because these mapped inside the supergene according to the previously defined chromosomal locations of the assembly scaffolds^14^ the supergene in the assembly by Yan *et al*. ^8^ (with blast, the genes align between position 8.39 and 8.40 Mb of chromosome 16 in GCA_009650705.1, whereas the supergene starts approximately at position 12.3 Mb). For each of the samples in the analysis, we created a consensus sequence for each of the genes, including coding sequence and introns, using bcftools consensus. We grouped the genes into non-overlapping windows of four concatenated neighboring genes. We used RAxML-NG^22^ (v0.9.0) with 10 random and 10 parsimony starting trees and the GTR+G model to build a maximum-likelihood tree of each window. We then performed phylogenetic topology weighting with Twisst^28^, using these trees as input, considering samples carrying SB and Sb as separate clades and *S. geminata* as the outgroup species.

The Twisst tool outputs a relative weight for each of 945 possible topologies for each window (given the seven clades in our analysis). We filtered our windows to keep those that supported a single topology with a relative weight greater than 0.9 (145 out of 212, or 68%) (Supplementary Fig. 13). Only eight out of 945 possible topologies were present in these 145 windows as the best topology. We classified these eight topologies by the evolutionary scenario that they support (Supplementary Fig. 14a). For the 39 windows mapping to the supergene region, the majority supported an introgression scenario (25, or 64%), of which 20 supported the introgression from *S. invicta/macdonaghi* to *S. richteri* (51% of the windows in the supergene) and the remaining 5 supported the opposite direction of introgression (13%; Supplementary Fig. 14b). Additionally, only 10 windows (26% of the windows in the supergene) supported the scenario of Sb as a trans-species polymorphism with ancestral origin. A further 3 windows (8%) supported the species tree topology. On the other hand, all the windows mapping to chromosome 1 (79 windows) and to the region outside the supergene in chromosome 16 (27 windows) supported the species tree.

We used blast to map the sequences of the breakpoints of the three previously identified inversions^8^ (In(16)1, In(16)2 and In(16)3) on the coordinates of the assembly used in our study (Supplementary Fig. 13). The largest inversion, In(16)3, covering 9.59 Mbp of assembled chromosome, included 47 windows, of which 37 (79%) had a single topology with a relative weight greater than 0.9. Of these, 18 (49%) supported the *S. invicta* to *S. richteri* introgression scenario, while 10 (27%) supported the trans-species ancestral polymorphism scenario. The inversion In(16)2 region of the supergene is 0.84 Mbp long and included 4 windows, of which 2 had a single topology with a weight greater than 0.9. Both supported the *S. invicta* to *S. richteri* introgression scenario. The inversion In(16)3 is 1.2 Mbp long, and included none of the windows in our analysis. We conclude that inversions 1 and 2, which together account for the vast majority of the assembled portion of the supergene, support the introgression scenario over the other scenarios.

#### Supplementary Note 8: Evidence that the recent introgression of Sb into *S. megergates* was mediated by *S. invicta/macdonaghi* queens

Above, we concluded that one of the introgressions of *S. megergates* (introgression 3 in Fig. 1e) occurred recently (Supplementary Notes 1 and 3). To explore whether this introgression could have been the result of the mating of a *S. invicta/macdonaghi* queen and a *S. megergates* male, we analyzed the mitochondrial haplotype carried by samples in the introgressed *S. megergates* colony and three *S. invicta*/*macdonaghi* colonies collected within 5.2 km. We found that all eight individuals we sampled from these colonies share the same mitochondrial haplotype (Supplementary Fig. 16). Because mitochondria are transmitted almost exclusively via queens, the most parsimonious explanation is that the current *S. megergates* lineage results from an original hybridization between a heterozygous (*SB/Sb*) *S. invicta/macdonaghi* queen and a *S. megergates* male, followed by backcrosses with *S. megergates* males. This would allow for the loss of *S. invicta/macdonaghi* nuclear haplotypes in the *S. megergates* lineage, despite the maintenance of the introgressed mitochondrial and Sb haplotype. The backcrossing between hybrids and *S. megergates* males could result from either pre-zygotic processes (*e.g.*, mating preference) or post-zygotic processes (*e.g.*, allelic incompatibilities reducing the fertility of hybrids).

In *Solenopsis,* an additional queen is accepted into a colony if both she and a large proportion of the workers in the colony carry the Sb haplotype^29^. We hypothesize that the original Sb-bearing hybrid joined an existing *S. invicta/macdonaghi* multiple-queen colony instead of founding an independent (single-queen) colony. This process would have allowed a hybrid queen to benefit from a social buffer against the possible negative effects of allelic incompatibilities in her genome.

The alternative mode of introgression, where Sb is transmitted by Sb-bearing *S. invicta/macdonaghi* males mating with *S. megergates* females, is less parsimonious, requiring an additional introgression of the mitochondrial haplotype from *S. invicta/macdonaghi* females to the hybrid lineage.

#### Supplementary Note 9: Restriction Fragment Length Polymorphism assay to identify the *Gp-9* genotype

We performed Restriction Fragment Length Polymorphism assays of phenol-chloroform DNA extraction of samples to identify their *Gp-9* genotype^30^. The *Gp-9* marker differentiates two allele types: the *Gp-9B­-*like alleles, associated with SB, and *Gp-9b-*like alleles, is associated with Sb. For the assay, we amplify a 828-bp fragment by PCR (95°C 3 min, 33 cycles of 95°C for 30s, 58°C for 30s and 72°C for 60s, 72°C for 5 min), using primers Gp-9_169.for (5' GGCCAGCACAAAACCAATC 3', Metabion) and Gp-9_490.rev (3' GTATGCCAGCTGTTTTTAATTGC 5', Metabion) in a 15 µl reaction (PeqLab high specificity buffer 1.8 µl, Peqlab Taq 0.05 µl, New England Biolabs dNTPs 0.3µl, forward primer 0.3 µl, reverse primer 0.3 µl, DNA 10 ng), followed by direct digestion by BsaAI (New England Biolabs, 0.3 µl) at 37°C for 4 h. The digested PCR products were visualized in a 1% Agarose gel with GelRed under UV light and fragments determined: the *Gp-9B*-like alleles yield two fragments (545 and 283 bp), whereas the *Gp-9b*-like alleles yield three (428, 283, and 117 bp).

We first used this assay on pools of 10 workers to identify the colony social form, and then on individual male samples to select samples carrying either SB or Sb.

#### Supplementary Note 10: Phylogenetic analysis of *Solenopsis* using mitochondrial sequences

We analyzed mitochondrial sequences from each individual in order to understand patterns of hybridization between divergent individuals and maternal transmission together with the supergene. Variants on the mitochondrion (*Solenopsis invicta* NC_014672.1) were called separately for each individual using freebayes^31^ v1.2.0 (--min-alternate-fraction 0.7 --min-alternate-total 3 --min-coverage 5 --use-best-n-alleles 1 --ploidy 1 --haplotype-length 20 --min-base-quality 25), followed by filtering low quality variants (Q<30), decomposition and filtering indels using vcflib (<https://github.com/vcflib/vcflib>). For each individual, a consensus FASTA based on its respective VCF file was extracted using bcftools^32^, and a multiple sequence alignment generated using MAFFT ^33^, including published fire ant mitochondrial sequences^34^ (*S. invicta*: NC_014672.1, HQ215540 and HQ215538; *S. richteri*: NC_014677; *S. geminata*: NC_014669). The sample gem1 was removed due to signs of reference bias in the variant detection and long-branch attraction. Using one partition per mitochondrial gene or non-coding region (see code on GitHub), the best substitution models were determined using ModelTest-NG^35^ v0.1.6 (parameters: -d nt -T raxml --rngseed 2 -t fixed-ml-gtr) and both used for maximum likelihood phylogenetic tree inference using RAxML-NG^22^ (parameters: --tree pars{50},rand{50} --seed 2 --blopt nr_safe --bs-trees 1000 --bs-metric fbp,tbe --bs-cutoff 0.05 --model model-per-partition.txt) with 100 starting trees (Supplementary Fig. 16). The bootstrapping converged after 150 bootstraps. The tree was rooted to *S. geminata* (NC_014669). For better visualization, the root taxon (*S. geminata*) was removed from Supplementary Fig. 16.

### Supplementary Figures

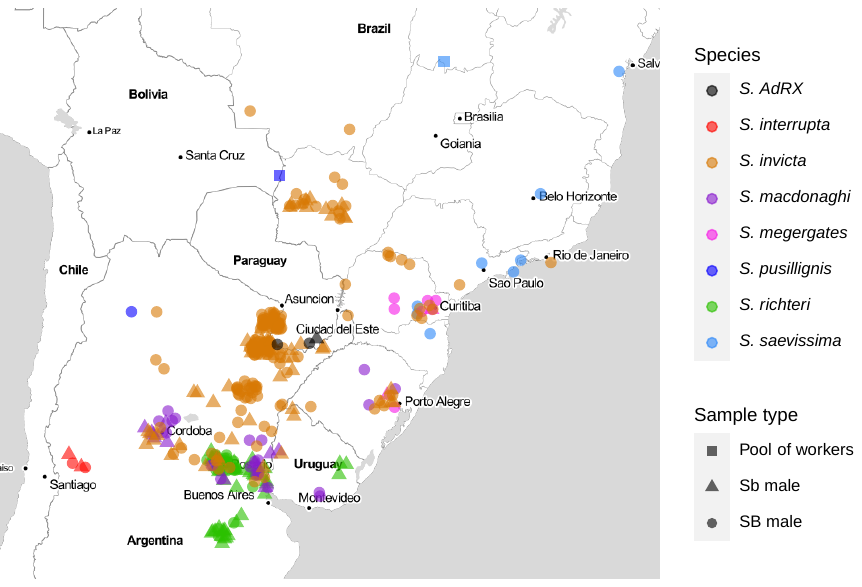

#### Supplementary Figure 1: Map of collection sites of the 349 South American samples

This represents 95% of the 368 samples used for this study (Supplementary Data 1). Color indicates species, and shape represents sample type (including supergene variants for males). Samples from French Guiana (n = 2), North America (n = 16) and Thailand (n = 1) are not shown. Locations are plotted with a slight jitter to help differentiate adjacent samples. This plot was built using the R package ggmap v3.0.0^36^, on map tiles by Stamen Design (under the CC BY 3.0 license, <http://creativecommons.org/licenses/by/3.0>) and map data by OpenStreetMap (under the ODbL license, <http://www.openstreetmap.org/copyright>).

#### Supplementary Figure 2: Dated maximum likelihood phylogenetic tree of fire ant individuals and species

An age calibrated phylogenetic tree was inferred, based on a concatenated gapless alignment of ~2.9 million nucleotides from 2,161 conserved single-copy ortholog coding sequences detected in individual genome assemblies outside the *S. invicta* chromosome 16, *i.e.*, chromosomes 1–15. Dating was calibrated by two node ages (*S. fugax*: -25.2 mya and *S. geminata*: -3.91 mya^5,6^). Blue bars display confidence intervals of inferred divergence times based on 100 replicates. Support for all nodes was 100% (1000 ultrafast bootstraps) and is not shown on the nodes, with the exception for two nodes (grey italic). Numbers on nodes show the inferred divergence times in bold for species’ divergence and non-bold for intraspecific divergence in million years.

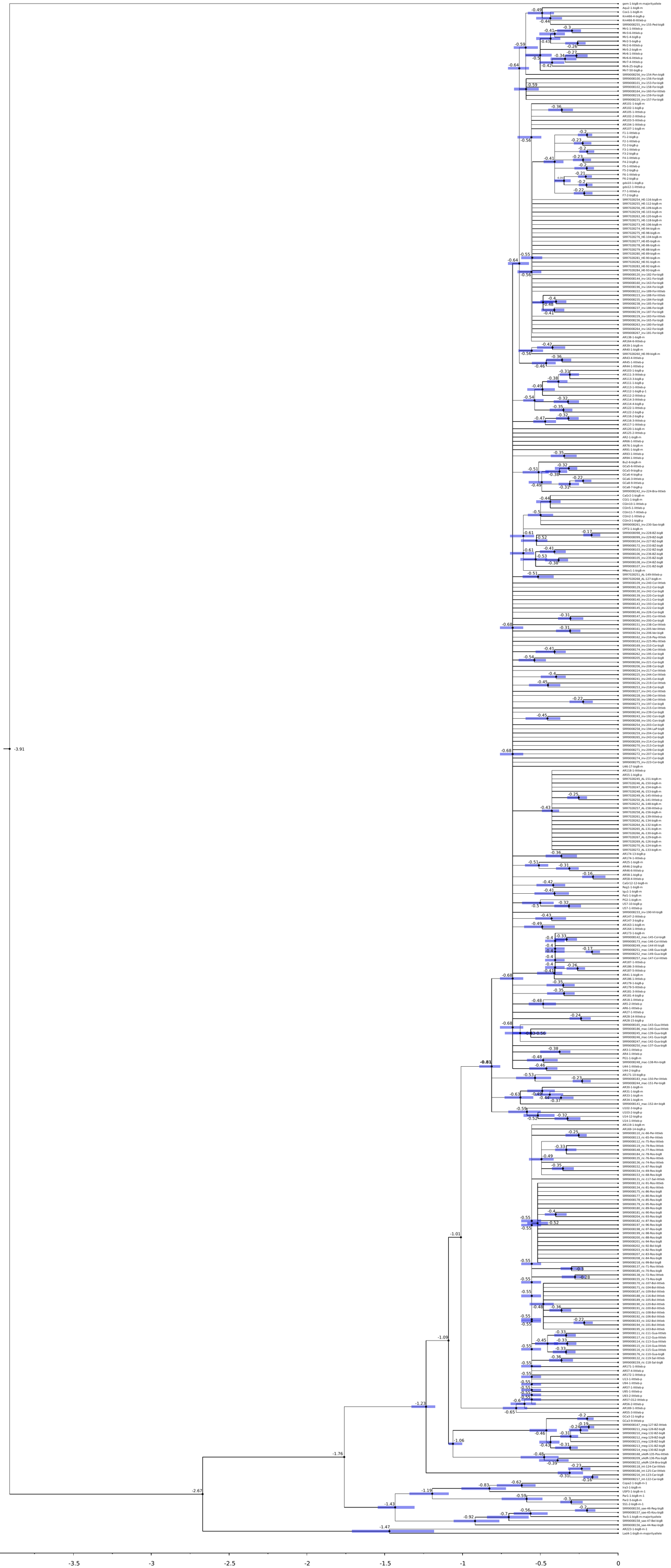

#### Supplementary Figure 3: Age calibrated phylogenetic tree of chromosomes 1–15

Tree was built with all samples present in Fig. 1e. Major nodes were calibrated with ages inferred in Supplementary Fig. 2. Bars display confidence intervals from 100 replicates.

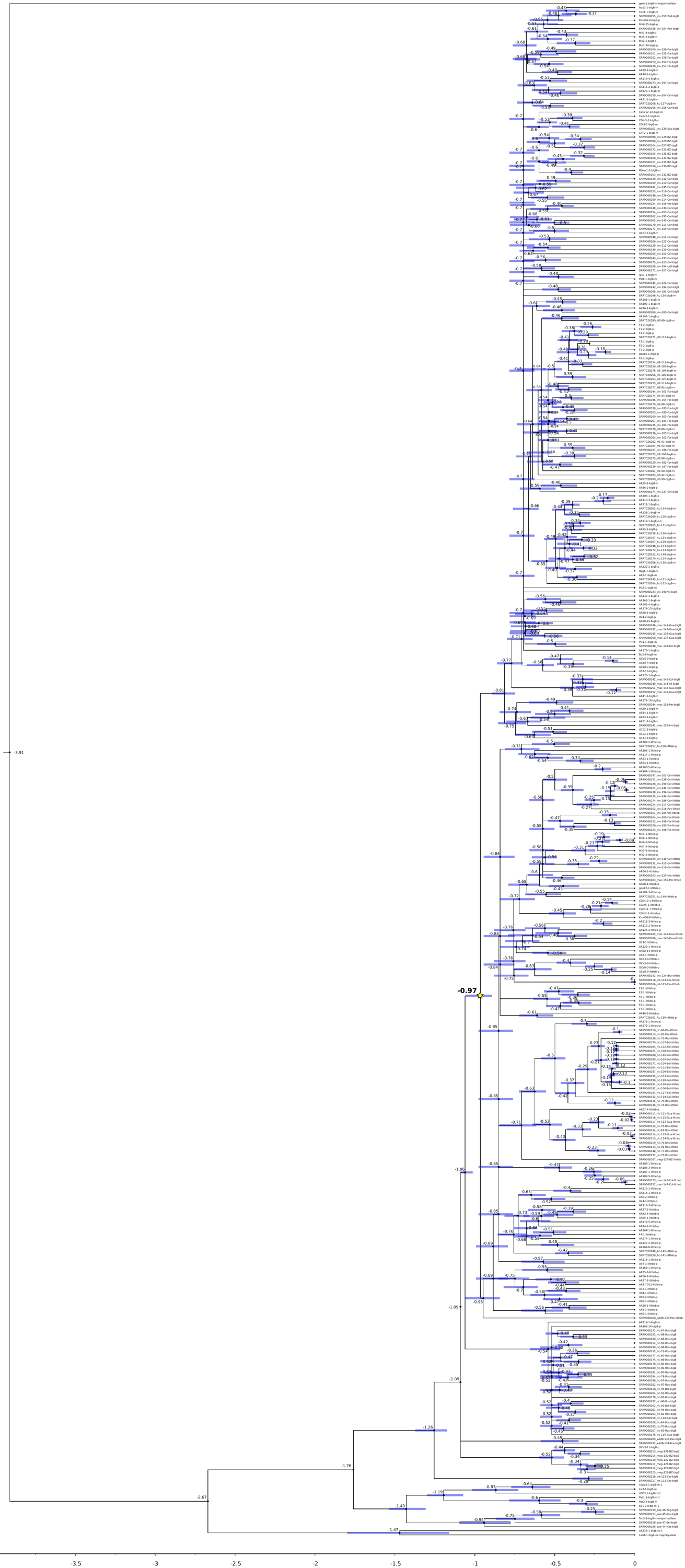

#### Supplementary Figure 4: Age calibrated phylogenetic tree of the supergene

Tree was built with all samples present in Fig. 1e. Major nodes were calibrated with ages inferred in Supplementary Fig. 2. Bars display confidence intervals from 100 replicates.

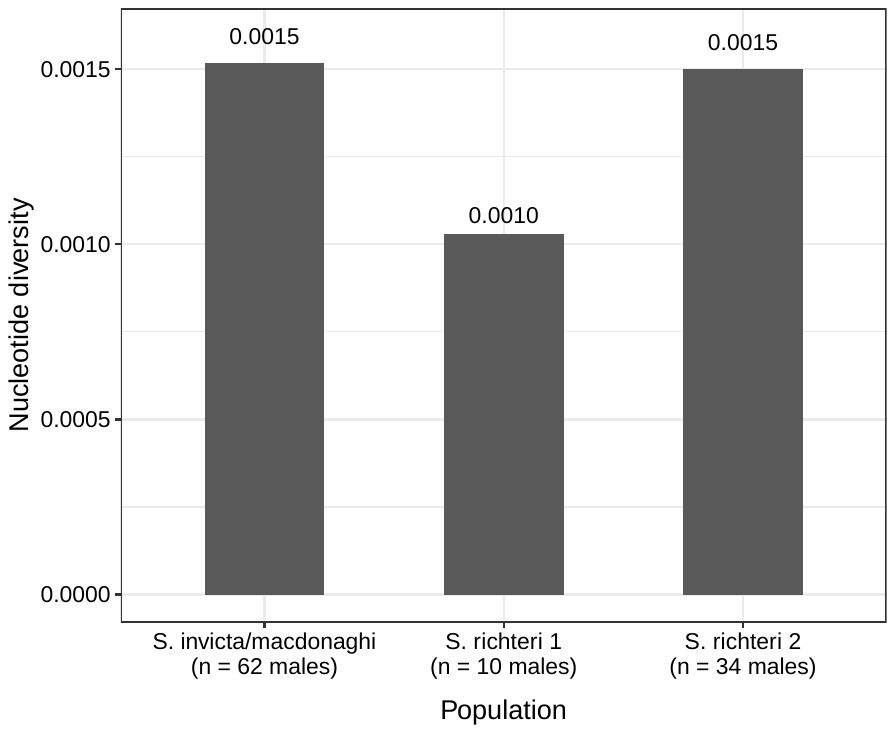

#### Supplementary Figure 5: Nucleotide diversity (π) of Sb in *S. invicta/macdonaghi* and each of the two introgressed clades of *S. richteri*

The *S. richteri* clades represent introgressions 1 and 2 in Fig. 1e. Nucleotide diversity (π) of Sb was calculated from 1 kb extended single-copy gene regions, summed across all regions and divided by their total size.

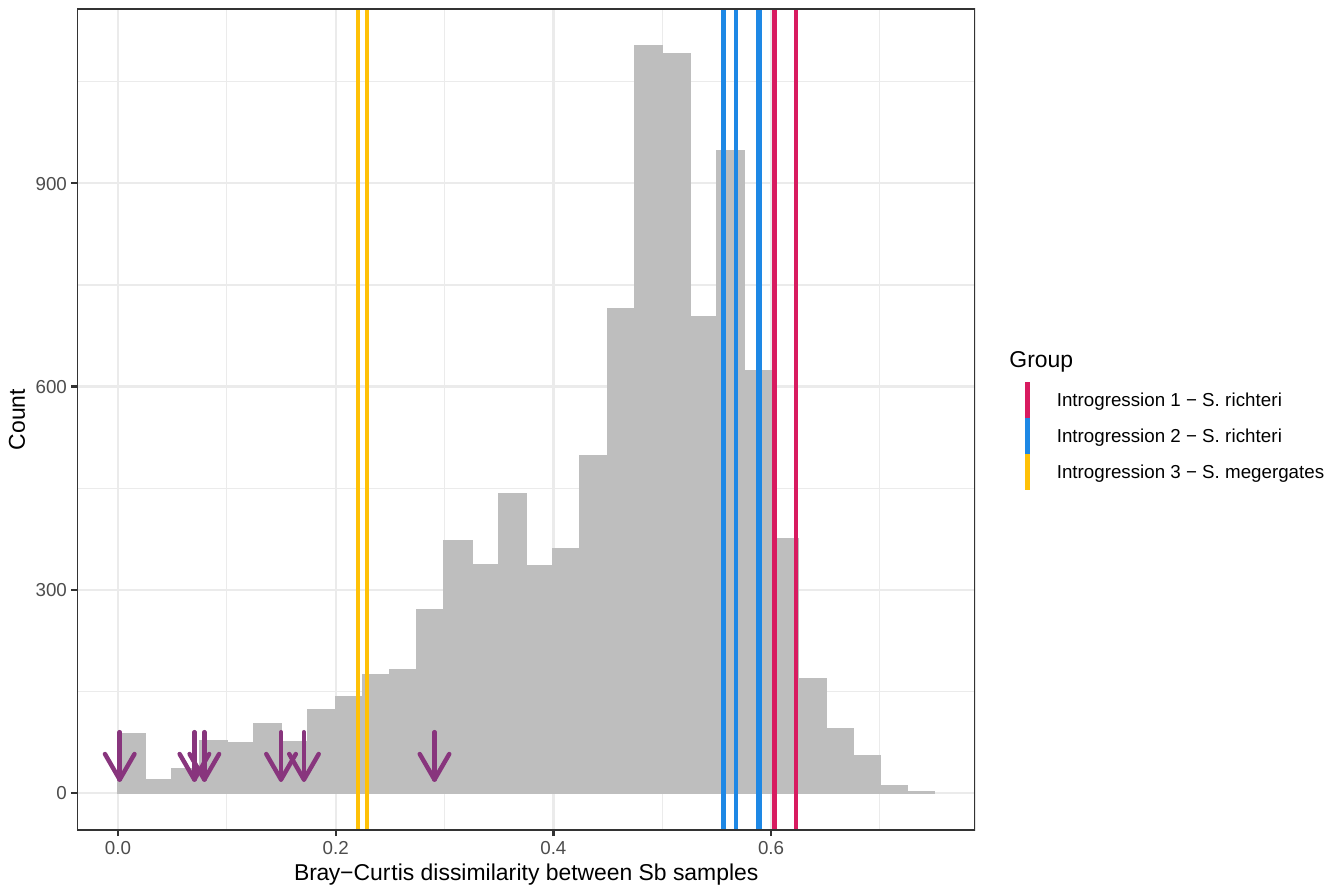

#### Supplementary Figure 6: Bray-Curtis dissimilarities between Sb haplotypes, highlighting the dissimilarity between introgressed haplotypes and respective outgroups

We measured the Bray-Curtis dissimilarity index with R package vegan^1^ between Sb haplotypes, based on the genotypes of the 24,435 SNPs that were bi-allelic among the Sb samples and mapped to the supergene (Supplementary Data 3). The distribution of dissimilarity values is shown in grey. The vertical lines represent:

- In red: The mean pairwise dissimilarity values between *S. richteri* samples in introgression 1 and the two closest *S. invicta/macdonaghi* samples (U57-1-littleb-p and AR118-1-littleb-p)
- In blue: The mean pairwise dissimilarity values between *S. richteri* samples in introgression 2 and the two closest *S. invicta/macdonaghi* samples (AR46-6-littleb-p, SRR7028257_AL-158-littleb-p, AR102-2-littleb-p and AR93-1-littleb-p)
- In orange: The pairwise dissimilarity values between the single *S. megergates* male (introgression 3) and each of the three *S. invicta/macdonaghi* Sb males in its clade (GCa5-6-littleb-p, GCa6-3-littleb-p and GCa8-9-littleb-p)

As a comparison, we also highlight the range of dissimilarities measured between Sb haplotypes of samples collected from within the same colony (purple arrows).

#### Supplementary Figure 7: Phylogenies of *Solenopsis* inferred with concatenated SNPs

Phylogenetic analyses of concatenated sequences support the pattern of Sb introgression.

(left) Phylogeny based on SNPs from chromosomes 1-15 (5,450 loci), with all sampled individuals grouping according to species.

(right) The phylogeny of SB and Sb haplotypes of chromosome 16 (210 loci) deviates from species-level monophyly, supporting the pattern of introgression of the Sb supergene variant from *S. invicta/macdonaghi* into other *Solenopsis* species (Fig. 1e). Nodes display transfer bootstrap expectation values greater than or equal to 80%. The scale bars indicate substitutions per site.

#### Supplementary Figure 8: Heatmap of average introgression fractions estimated using 50 samples

We used QuIBL to test for introgression based on the internal branch length distributions of sample triplets. The quiblR function “getIntrogressionSummary” evaluates all triplets discordant with the species tree and averages the value "mixprop2*count/total trees'' for each pair of samples (mixprop2 corresponds to the non-ILS component of the inferred distribution). Color labels along axes correspond to different *Solenopsis* species.

#### Supplementary Figure 9: Branch length distributions for triplets of interest.

We extracted the internal branch lengths of triplet gene trees using the quiblR function "getPerLocusStats". The distribution of internal branch lengths reveals two peaks, indicative of introgression.

#### Supplementary Figure 10: Heatmap of average introgression fractions estimated using 17 samples

We used QuIBL to test for introgression based on the internal branch length distributions of sample triplets. The quiblR function “getIntrogressionSummary” evaluates all triplets discordant with the species tree and averages the value "mixprop2*count/total trees'' for each pair of samples (mixprop2 corresponds to the non-ILS component of the inferred distribution). Color labels along axes correspond to different *Solenopsis* species.

#### Supplementary Figure 11: Branch length distributions for triplets of interest

We extracted the internal branch lengths of triplet gene trees using the quiblR function "getPerLocusStats". The distribution of internal branch lengths reveals two peaks, indicative of introgression.

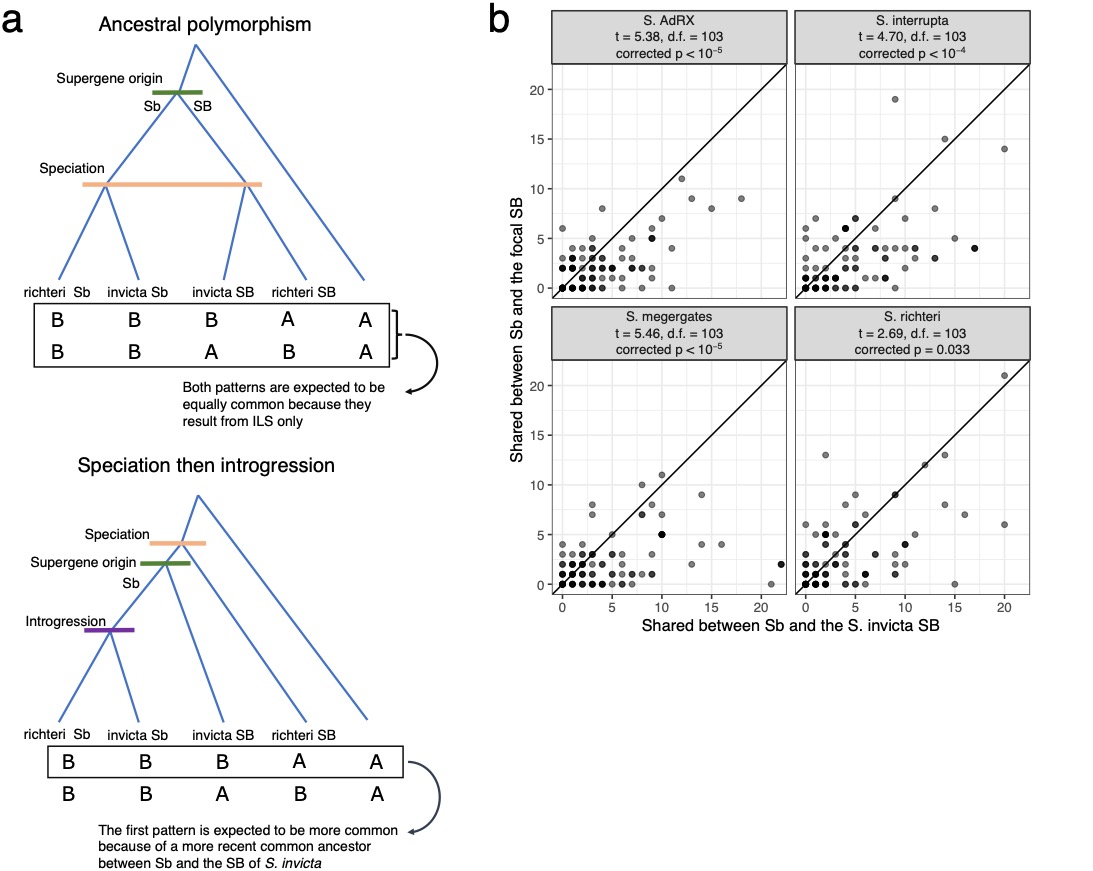

#### Supplementary Figure 12: Derived alleles shared by Sb and the SB variant of either *S. invicta* or each of four other species that carry the supergene (BBBAA-BBABA test)

**a.** Expected pattern under the ancestral polymorphism scenario (top) or the introgression scenario (from *S. invicta/macdonaghi* to the focal species, bottom), using the example of *S. richteri* as the focal species.

**b.** BBBAA-BBABA test including the early diverging species. We performed four comparisons, each time taking two *S. invicta* samples (one SB and one Sb) and two samples from another species (SB and Sb), as well as a sample representing the outgroup species *S. saevissima.*

For each of 104 genes mapped to the supergene, we counted the number of alleles carried by the two Sb individuals in each comparison and the SB individual of either *S. invicta* or the focal species; there were more of such shared alleles in the *S. invicta* SB than there were for the SB of the other species (two-tail paired t-test, Bonferroni-corrected *p* < 0.05; test parameters given in each panel, degrees of freedom denoted as “d.f.”).

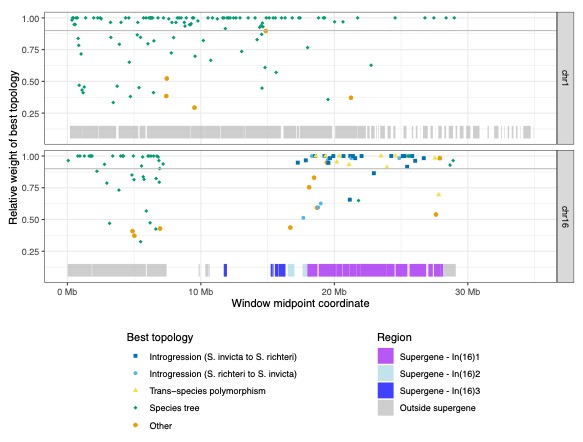

#### Supplementary Figure 13: Evolutionary scenario represented by the topology with highest support in the phylogenetic weighting analysis of non-overlapping windows

Relative weight of the best supported topology in the phylogenetic weighting analysis (Twisst^28^) of non-overlapping 213 windows of four concatenated genes, colored by the evolutionary scenario (see Supplementary Fig. 14 for the most highly supported topologies). The bars at the bottom of each panel represent the chromosomal location of the assembly scaffolds, as previously inferred^14^; these are colored according to whether they are affected by the inversions identified by Yan *et al*.^8^, which we mapped by aligning the protein-coding genes closest to the breakpoint sequences to the genome assembly used here. No window had a midpoint within In(16)3. Four windows had a midpoint within In(16)2, of which two had a topology with relative weight greater than 0.9, both supporting the scenario of introgression between *S. invicta* and *S. richteri*.

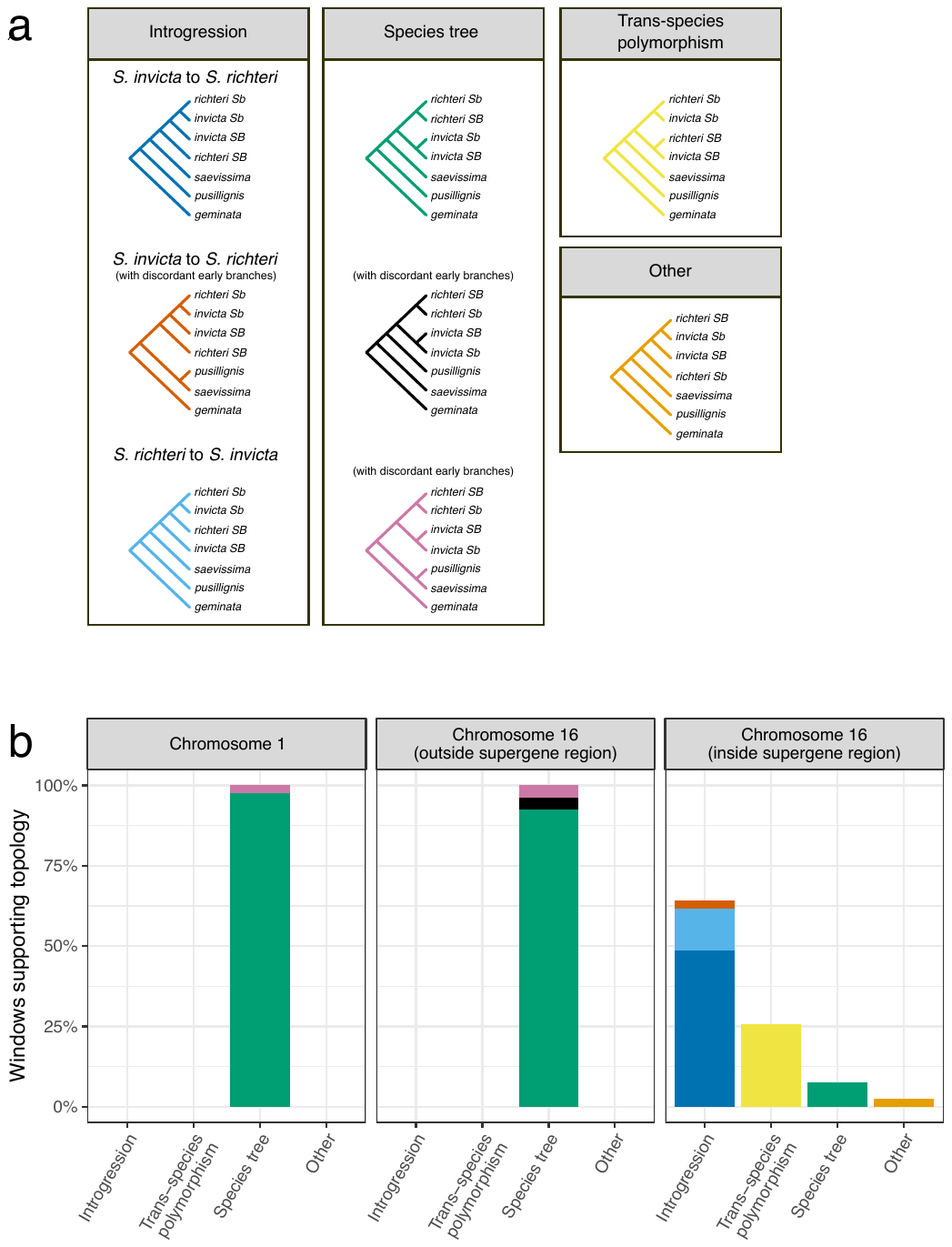

#### Supplementary Figure 14: Phylogenetic weighting analysis of non-overlapping windows represented regions inside and outside the supergene

**a.** Best topologies for each of the 145 windows (out of 212, or 68%) that have a single topology with a relative weight greater than 0.9. The topologies are grouped into boxes depending on the evolutionary scenario that they represent.

**b.** The evolutionary scenario supported by the best topology of the 79 windows in chromosome 1, the 27 windows outside the supergene region of chromosome 16 and the 39 windows inside the supergene region of chromosome 16. As in panel a, we only considered the 14 windows that have a single topology with a relative weight greater than 0.9; topologies are colored according to panel a. Note that, for the windows in the supergene region, the “Species tree” scenario would suggest an independent origin of Sb in each of the species.

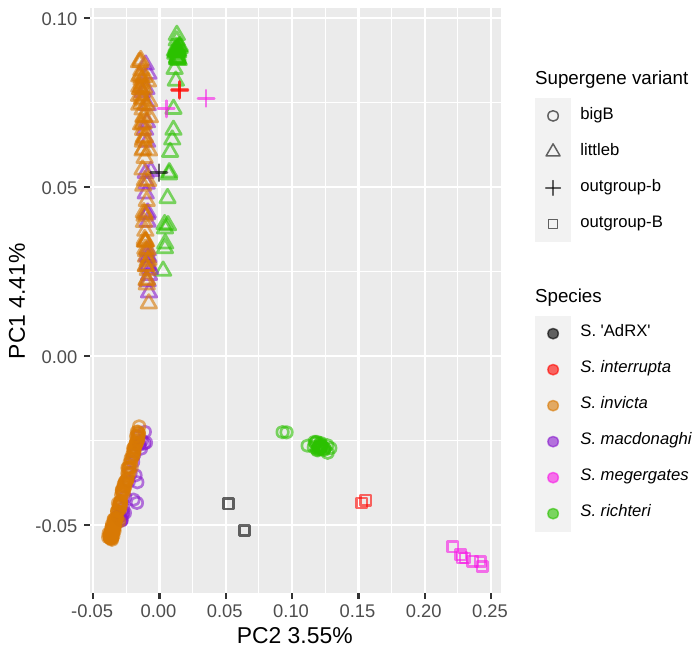

#### Supplementary Figure 15: Principal component analysis of supergene SNP data

Data points represent 354 males of *S. invicta/macdonaghi*, *S. richteri* and the closely related species *S. megergates*, *S. AdRX* and *S. interrupta*.

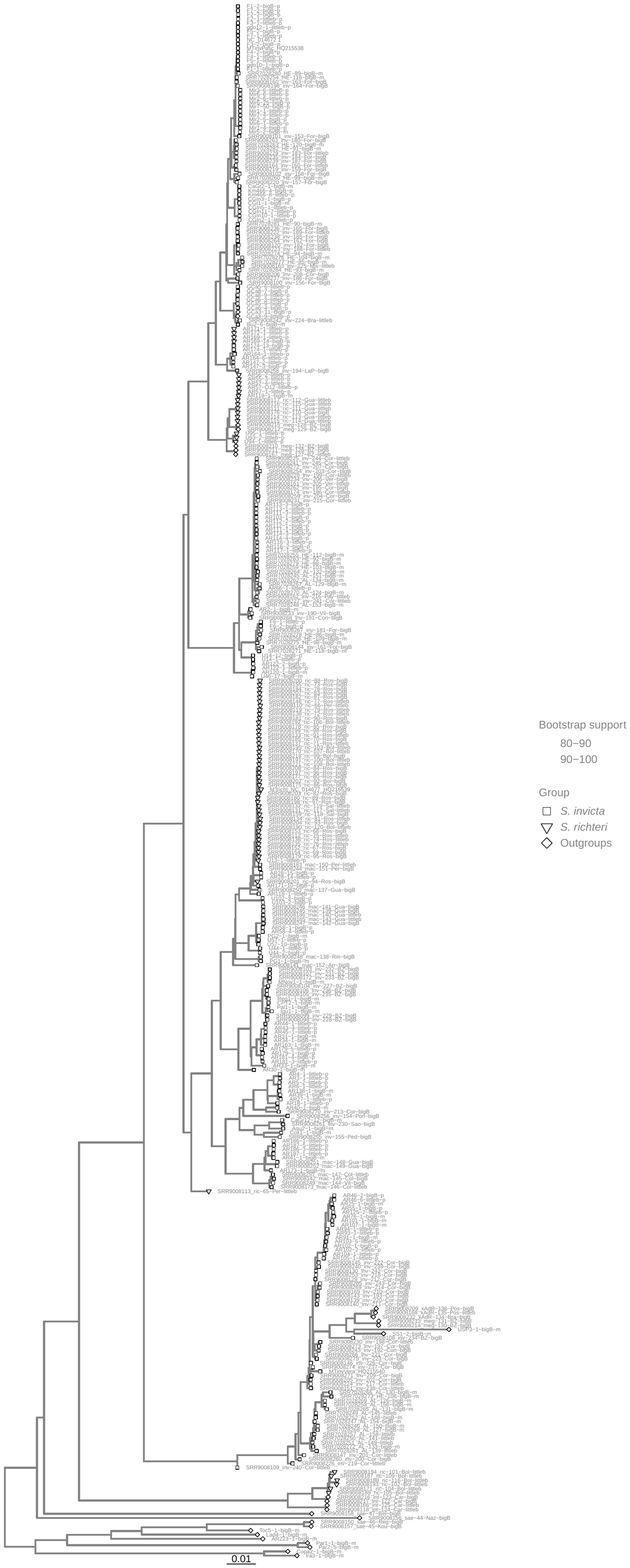

#### Supplementary Figure 16. Phylogeny of *Solenopsis* inferred with mitochondrial sequences

Maximum likelihood phylogenetic tree of *Solenopsis* inferred from full mitochondrial sequences of 367 non-*geminata* samples used in this study and five previously published full mitochondrial sequences (see Supplementary Note 10). We ran RAxML-NG with one partition and model per gene (50 random and 50 parsimony starting trees, 150 bootstraps). The tree was rooted to *S. geminata* (NC_014669), although for better visualization, this taxon was removed from the figure.
